# Hierarchically Vascularized and Implantable Tissue Constructs created through Angiogenesis from Tissue-Engineered Vascular Grafts

**DOI:** 10.1101/2024.04.29.591796

**Authors:** Hazem Alkazemi, Geraldine M. Mitchell, Zerina Lokmic-Tomkins, Daniel E. Heath, Andrea J. O’Connor

## Abstract

A major roadblock in implementing engineered tissues clinically lies in their limited vascularization. After implantation, such tissues do not integrate with the host’s circulation as quickly as needed, commonly resulting in loss of viability and functionality. This study presents a solution to the vascularization problem that could enable the survival and function of large, transplantable, and vascularized engineered tissues. The technique allows vascularization of a cell laden hydrogel through angiogenesis from a suturable tissue-engineered vascular graft (TEVG) constructed from electrospun polycaprolactone with macropores. The graft is surrounded by a layer of cell-laden gelatin-methacryloyl hydrogel. The constructs are suturable and possess mechanical properties like native vessels. Angiogenesis occurs through the pores in the graft, resulting in a hydrogel tcontaining an extensive vascular network that is connected to an implantable TEVG. The size of the engineered tissue and the degree of vascularization can be increased by adding multiple TEVGs into a single construct. The engineered tissue has the potential to be immediately perfused by the patient’s blood upon surgical anastomosis to host vessels, enabling survival of implanted cells. These findings provide a meaningful step to address the longstanding problem of fabricating suturable pre-vascularized tissues which could survive upon implantation *in vivo*.

## 1. Introduction

Vascularization is a major challenge that limits the clinical implementation of tissue engineering [1, 2]. Specifically, most three-dimensional, engineered tissues exhibit poor survival upon implantation, largely due to oxygen transport limitations. *In vivo*, most tissues require blood to be supplied via a capillary network since sufficient oxygen levels can only diffuse ∼200 μm from blood vessels in cell-dense tissues [3, 4]. In contrast, engineered tissues are not rapidly connected to the patient’s vascular supply upon implantation. Even if an engineered tissue is generated to contain a capillary network *in vitro* (referred to as ‘pre-vascularization’ hereafter), its *in vivo* survival will depend on the slow process of inosculation (functional unification of donor and host capillary networks). This process takes days to weeks before the network is well connected to and perfused by the recipient’s circulating blood, and the diffusion-based transport of oxygen during this time is not sufficient to maintain viability and function of the engineered tissue [5, 6]. Novel technologies are required to enable the rapid perfusion of engineered tissue by the host blood supply.

Significant research has occurred over the last 30 years to address the vascularization problem. For instance, researchers have developed various approaches to pre-vascularize engineered tissues prior to implantation, thereby shortening the time needed for inosculation [7-9]. Additionally, tissue engineered constructs with porous or channels may be kept alive *in vitro* using perfusion bioreactors that circulate oxygenated medium. Examples of these approaches include microfluidic systems, sacrificial molding, and many others [10-12]. For instance, Kolesky, et al. fabricated large tissues (>1 cm thickness) *in vitro* using 3D bioprinting techniques [13]. Despite these recent advances, engineered tissues still cannot be directly connected to the patient’s blood supply upon implantation resulting in poor survival of the engineered tissue and limiting the clinical success of potentially lifesaving tissue engineering strategies [14].

A pre-vascularized, engineered tissue requires immediate connection to the body’s circulation to ensure survival and function of the new tissue by enabling rapid delivery of sufficient oxygen and nutrients [15]. Immediate connection can only be achieved if the engineered tissue incorporates a medium sized artery and vein continuous with the capillary circulation. These larger vessels can be directly anastomosed to host vessels of similar size to ensure rapid blood perfusion of implanted engineered tissues. However, many engineered vessels and tissues are fabricated using soft hydrogels, and these materials lack the mechanical strength, suturability, and adequate burst pressure to be directly connected to the patient’s circulation [16].

Plastic and reconstructive surgeons often use free tissue flap procedures to replace soft tissue, such as fat for breast reconstruction after mastectomy [17]. In the free flap procedure following mastectomy, a piece of vascularized fat – including its supplying artery and vein – is removed from a donor site on the patient’s body. The donor tissue’s artery and vein are connected to recipient vessels in the chest wall to allow immediate blood perfusion of the tissue, and the tissue is then implanted to restore volume and contour in place of the removed breast tissue [18]. This procedure illustrates the need for a hierarchical and suturable blood vessel network within a tissue to enable survival upon implantation, and it provides an attractive template for how to fabricate engineered tissues that can be immediately perfused by the recipient’s blood upon implantation.

We aimed to overcome the current limitations of vascularization in tissue engineering by fabricating a hierarchically vascularized, engineered tissue that is built around a suturable tissue-engineered vascular graft (TEVG), where the vascularization is grown via sprouting angiogenesis from the TEVG. Such a system would enable immediate perfusion of the engineered tissue *in vivo* by suturing the vascular graft to a patient’s macrovessels during implantation. Specifically, we aimed to fabricate a cell-laded hydrogel around a TEVG and to vascularize the hydrogel through angiogenesis from the macrovessel. Ideally, the fabrication process should be tunable to accommodate different cell types and customizable in terms of size and shape so that various tissues could be fabricated.

Recently, we developed TEVGs that mimic the biological architecture and key mechanical properties of native blood vessels (NBVs), including appropriate suture retention strength and burst pressure, to allow implantation. Additionally, these grafts have the advantage that they could be fabricated using a rapid and scalable process, facilitating translation and clinical implementation [19]. The vascular grafts are built on an electrospun tube of polycaprolactone (PCL) nanofibers that provide the necessary mechanical strength. PCL was chosen because it is a well-established biodegradable biomaterial that is conducive to fabrication via electrospinning. The fibers of the tubes are partially aligned in the axial direction through a freezing-induced alignment process, which results in an axially aligned and confluent endothelium after seeding with endothelial cells (EC), similar to that observed in NBVs [19]. The PCL layer is surrounded by a cast layer of gelatin methacryloyl (GelMA) laden with vascular smooth muscle cells (SMCs). GelMA was selected because it is a widely used biomaterial with a documented track record of biocompatibility and tunable mechanical and cell adhesion properties [20, 21]. By appropriately tuning the stiffness of the GelMA, we achieved spontaneous and rapid circumferential alignment and elongation of the SMCs, mirroring the characteristics observed in the media of NBVs [19].

No angiogenesis was observed from the lumen of these TEVGs into the surrounding GelMA layer. The lack of EC migration is likely due to the dense, fibrous electrospun layer, consistent with previous studies. [22, 23]. Previous research illustrates that 3-10 µm diameter pores are required for individual cell migration, and much larger pores – of the order of 100 µm diameter – are required to allow sustained capillary growth [24-26]. Similarly, in the process of sprouting angiogenesis from native blood vessels, several synchronized biological processes result in the formation of larger pores through EC basement membrane degradation, enabling the outgrowth of new vasculature [27]. In our TEVGs, the dense electrospun layer acts similarly to the intact basement membrane, by enabling the formation of a confluent endothelium, but preventing migration of ECs from the lumen and preventing sprouting due to the absence of pores of sizes conducive to angiogenic migration. In this contribution, we test the hypothesis that machining pores with sufficient diameter through the electrospun layer will enable EC migration from the lumen and vascularization of the hydrogel layer with capillary-like structures that are connected to the suturable macrovessel. To test this hypothesis, we created 150 µm diameter pores in the electrospun PCL tube by micro-machining prior to casting the GelMA layer. The large pore size should enhance EC migration from the lumen and infiltration into the hydrogel layer, key steps for angiogenesis initiation. This size of pores was also selected as it could foster the maturation of the new vasculature towards larger, arteriole- and venule-like vessels [28]. The results illustrated that macropores did facilitate EC migration from the lumen, and that appropriately controlling the stiffness of the GelMA hydrogel enabled the formation of an interconnected blood vessel network. We further hypothesized that using two endothelized vascular grafts would allow for the creation of a larger vascularized construct by providing two sources of ECs and shorter distances for angiogenesis, leading to the formation of a more extensive vascular network throughout the construct. This hypothesis was also supported by the data. This construct design provided two central medium sized vessels – similar to a vascular pedicle (adjacent artery and vein). This approach of combining pre-vascularized tissues with mechanically robust and suturable TEVGs to produce vascularized tissues has the potential to remove a longstanding challenge that has limited the field of tissue engineering since its inception and further unlock the as yet unmet potential of the field.

## 2. Results and Discussion

### 2.1. Tissue-Engineered Vascular Grafts with Pores show Comparable Mechanical Properties to Native Blood Vessels

The fabrication of the engineered tissue is schematically shown in **Figure 1**. The electrospun PCL scaffold (6 cm length, 2.5 mm inner diameter, 200 μm wall thickness) was prepared as previously described (**Figure 1a-i**) [19]. The graft’s electrospun layer was then machined to include 150 µm diameter pores with 1.5 mm spacing, as depicted in **Figure 1a-ii**. This processing resulted in tubes with evenly spaced pores, and SEM imaging showed that the pores pierce the PCL layer without significant damage to the electrospun scaffold (**Figure 1a-iii**). **Figure 1b** illustrates that a GelMA layer (either with or without SMCs) was cast and cured around the electrospun scaffold, the lumen was seeded with ECs to allow formation of an endothelial layer on the interior surface of the graft and the growth of a capillary network through the holes of the graft. In our previous work, the thickness of the GelMA layer was found to be approximately 800 µm using SEM analysis of freeze-dried samples [19]. The degree of methacrylation of the GelMA was quantified via ^1^H NMR spectroscopy by comparing the lysine peaks in the spectra of GelMA and gelatin as previously described [29]. The ^1^H NMR spectrum of gelatin and GelMA is shown in **Figure S1**. The degree of methacrylation of GelMA was found to be 72%, which ensures that the GelMA has sufficient mechanical strength after photo-crosslinking, while retaining the thermoresponsive properties of gelatin [30]. Finally, **Figure 1c** illustrates that multiple grafts can be used together to create larger constructs and to produce a more extensive capillary network.

**Figure 1.**
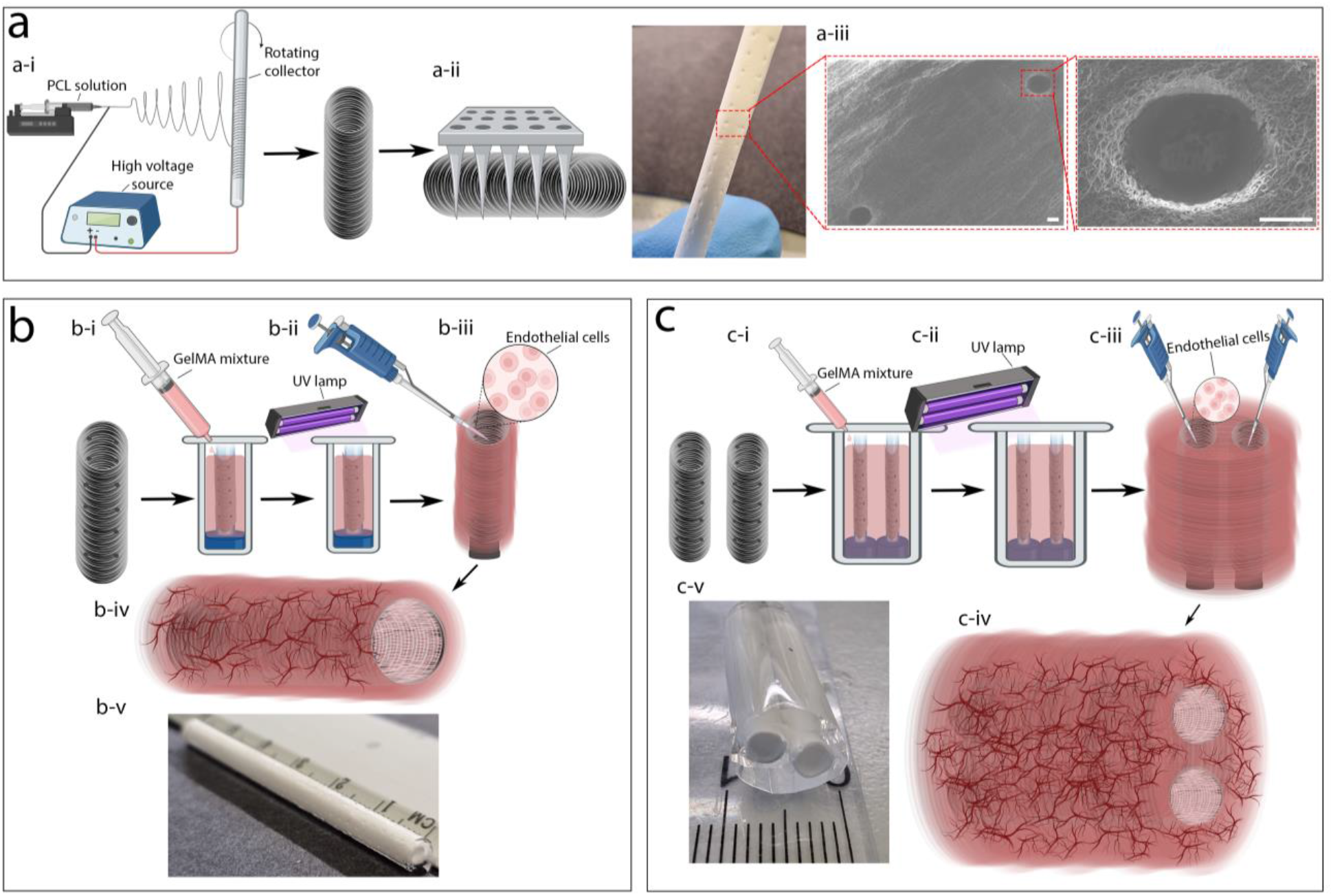
The process of fabricating engineered tissue. a-i) Tubular PCL scaffolds were fabricated via electrospinning and a-ii) machined to create macropores to enable angiogenesis. a-iii) The presence of the pores was confirmed through visual inspection and SEM imaging (scale bar: 100 µm). b-i and ii) A GelMA hydrogel layer (with or without SMCs) was cast and cured around the electrospun scaffold, and b-iii and iv) the luminal surface was seeded with endothelial cells to enable the formation of an endothelium and sprouting angiogenesis. b-v) Photograph of a TEGV with macropores. c) Creating a large, vascularized tissue by using two vascular grafts is also possible by c-i) casting a GelMA mixture around two electrospun tubes, c-ii) cross-linking the GelMA layer, and c-iii) Seeding ECs in the lumens of the PCL tubes. c-iv) This has the potential to generate large, vascularized tissues. c-v) Photograph of a large construct incorporating two TEVGs.

To ensure the mechanical stability of the vascularized tissue for clinical applications, we engineered the TEVGs to possess similar mechanical properties to the NBVs (saphenous veins and internal mammary arteries) commonly used in coronary artery bypass surgery, as reported in our previous study [19]. However, the machined pores could have reduced the mechanical properties of the grafts so we measured the burst pressure, suture retention strength, and tensile properties of TEVGs with and without the machined pores, as shown in **Figure 2**. The measured values were plotted next to literature-reported values for NBVs [31, 32], although the values were not compared by statistical tests due to differences in equipment and procedures used for data collection. As shown in **Figure 2a**, the burst pressure of the PCL tubes decreased a little due to the presence of the pores (p < 0.05), but it remained comparable to that of NBVs. The pores did not significantly impact the suture retention strength (p = 0.113), ultimate tensile strength (p = 0.335), or Young’s modulus (p = 0.262) of the vascular grafts, which were all comparable to those of NBVs (**Figure 2b-d**). Therefore, the vascular grafts could serve as a suitable scaffold for implanting the vascularized tissue *in vivo*. Moreover, the mechanical properties of the TEVGs are more similar to NBVs than currently used synthetic grafts that are significantly stiffer. The stiffness mismatch between NBVs and synthetic grafts is hypothesized to be a major contributor to synthetic graft failure by causing abnormal blood flow patterns, illustrating another advantage of the TEVGs developed here [33].

**Figure 2.**
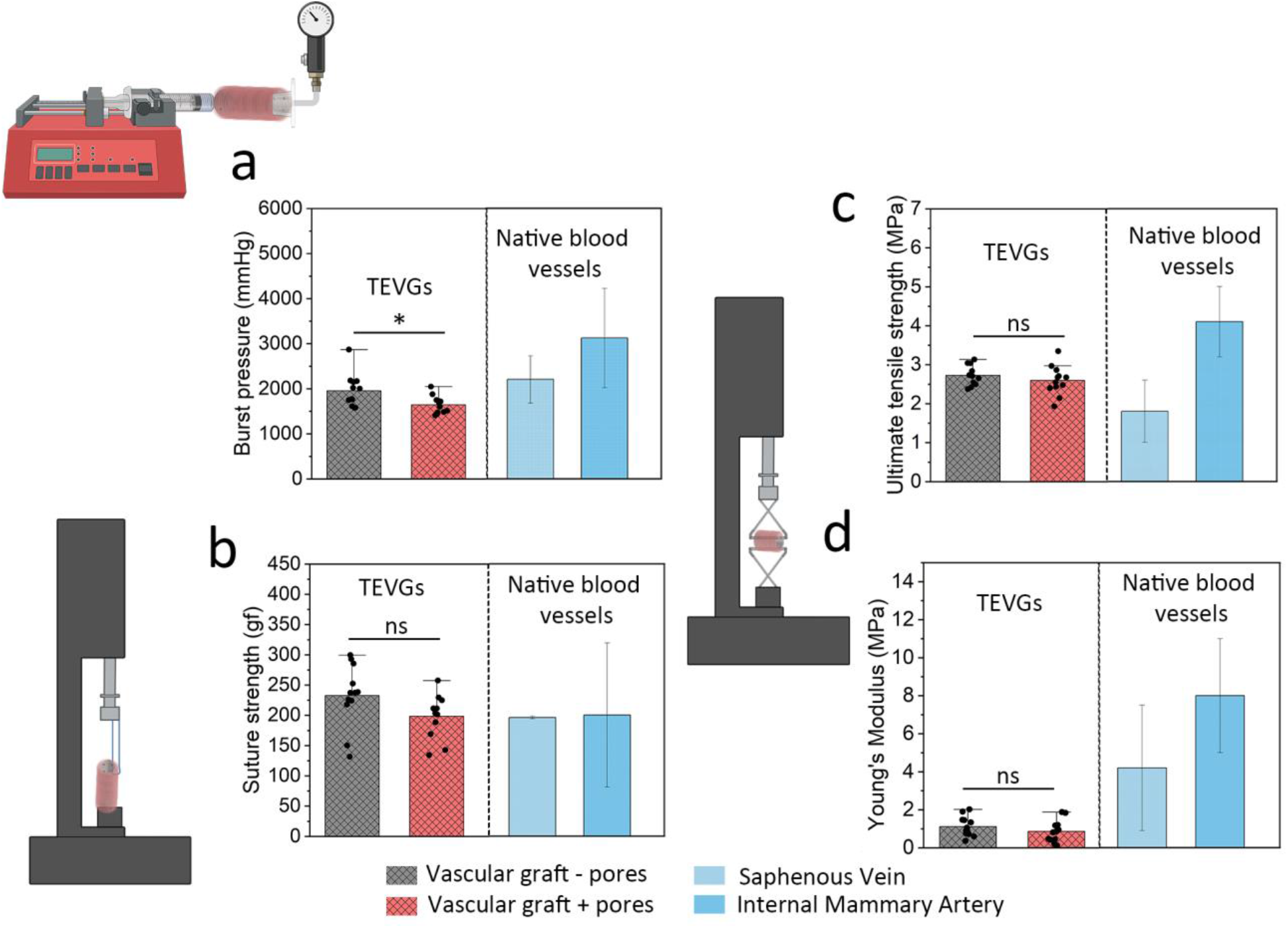
Mechanical properties of the vascular grafts before and after micro-machining compared to values of saphenous veins and internal mammary arteries from the literature [31, 32]. a) Burst pressure test results show the maximum pressure before bursting. b) Suture retention test results showing the maximum suture strength before failure. Tensile test results were analyzed to determine c) the ultimate tensile strength and d) the Young’s Modulus. Statistical analysis was performed to investigate the impact of micro-machining on the mechanical properties of the vascular grafts. Error bars represent the standard deviation (n = 12; *: p<0.05; ns: no significant difference, t-test). No statistical comparison was made between the vascular grafts and the NBVs since the data from the literature was collected using different methods and equipment.

### 2.2. Endothelial Cells Rapidly Form a Confluent Endothelium on the Vascular Graft Luminal Surface

NBVs have a confluent endothelial layer that confers anti-thrombogenic properties and regulates the selective permeability of molecules [34]. The ECs also participate in angiogenesis by migrating from the endothelium into the surrounding tissue in response to hypoxia signals [35]. Therefore, the vascular grafts should facilitate the formation of a confluent endothelium. To evaluate the endothelialization of the lumen after seeding the ECs, the cells were fixed and the samples were sectioned and imaged at different time points (days 1, 3, 5, 7, 10, and 14). **Figure 3a** shows an image of the cross section of the graft after 7 days, and it is observed that the entire circumference of the lumen has been covered by ECs. The confocal images of the endothelium at selected time points (**Figure 3b**) reveal proliferation of the ECs after seeding, and a confluent endothelium with complete coverage of the luminal surface with cells closely positioned next to one another appears to be established by day 5, as indicated by the CD31 staining. The analysis of the confocal images (**Figure 3c**) demonstrates that the endothelial coverage was minimal at day 1 (17±8%) but increased significantly at days 3 (40±14%) and 5 (87±5%) and remained at the same value for days 5 to 14 (91±3%). This saturation of the surface with ECs also illustrates the formation of a full endothelium. This rapid endothelialization of the lumen is beneficial for the quick fabrication of vascularized tissue, and it provides a source of ECs that can migrate into the surrounding hydrogel to form the capillary network.

**Figure 3.**
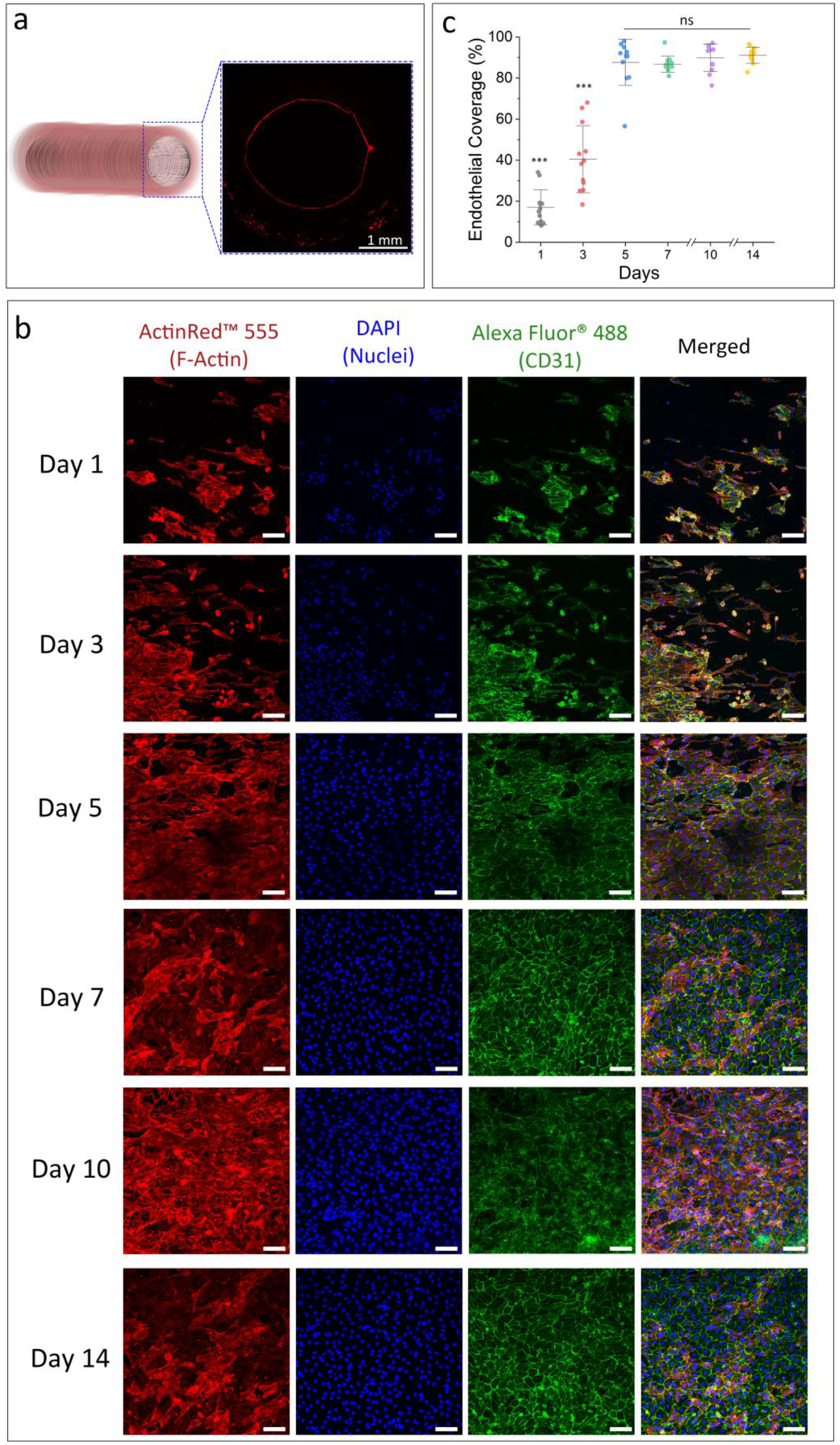
Endothelialization of the vascular grafts. a) A confocal image showing circumferential coverage of ECs on the lumen surface of the vascular graft after 7 days of culture, red: actin filaments stain (ActinRed™ 555), blue: nuclei stain (DAPI). b) Confocal images of ECs on the surface of the graft lumen at different time points showing the increase of endothelial coverage and proximity of the cells; scale bar = 50 μm. c) Quantitative analysis of EC coverage by area of the lumen over time. Error bars represent the standard deviation (n = 12; ***: p < 0.001, one-way ANOVA).

### 2.3. Micro-machined Pores Allow ECs to Migrate from the Lumen Toward the Surrounding Tissue

We hypothesized that micro-machining 150μm diameter pores into the vascular graft would enable sprouting angiogenesis and the growth of a capillary network into the surrounding hydrogel. **Figure 4a** shows the collective migration of ECs from the lumen of the PCL tube into the GelMA layer. This migration starts shortly after seeding the ECs in the lumen and increases with time as the endothelial coverage reaches confluence (∼90%) and the ECs have limited space for proliferation. **Figure 4b** qualitatively illustrates that the presence of the pores is required for significant migration of ECs from the lumen of the graft into the hydrogel. Without pores, few ECs are seen in the hydrogel layer, even after 7 days of culture. In contrast, many ECs are observed in the hydrogel layer in scaffolds that were micro-machined with the 150μm diameter pores. **Figure 4c** quantifies the number of cells within the GelMA layer at various timepoints post-seeding with ECs, and the quantitative data supports the qualitative observation from the confocal images. Specifically, engineered tissues without pores have few ECs in the GelMA layer, likely due to the dense PCL fiber network hindering the passage of the cells. This result was expected as previous literature has reported that ECs require a minimum of 3-10 µm diameter continuous pores to allow migration [24, 25]. In contrast, a statistically significant increase (p < 0.001) in the number of cells was observed for samples with pores, and the number of cells in the GelMA layer increased over time during culture. These data highlight the importance of macropores for the migration of a large number of ECs from the lumen of the graft into the hydrogel layer. This migration behavior is crucial for the formation of a vascular network in GelMA and its connection with the vascular graft to support the delivery of oxygen and nutrients to the engineered tissue. In this study, one pore geometry was used (150 µm pores with 1.5 mm spacing) to illustrate that the pores enabled endothelial cell migration into the GelMA layer and network formation. In future work, we will vary this spacing to optimize the endothelial network formation in the gel while maintaining graft integrity.

**Figure 4.**
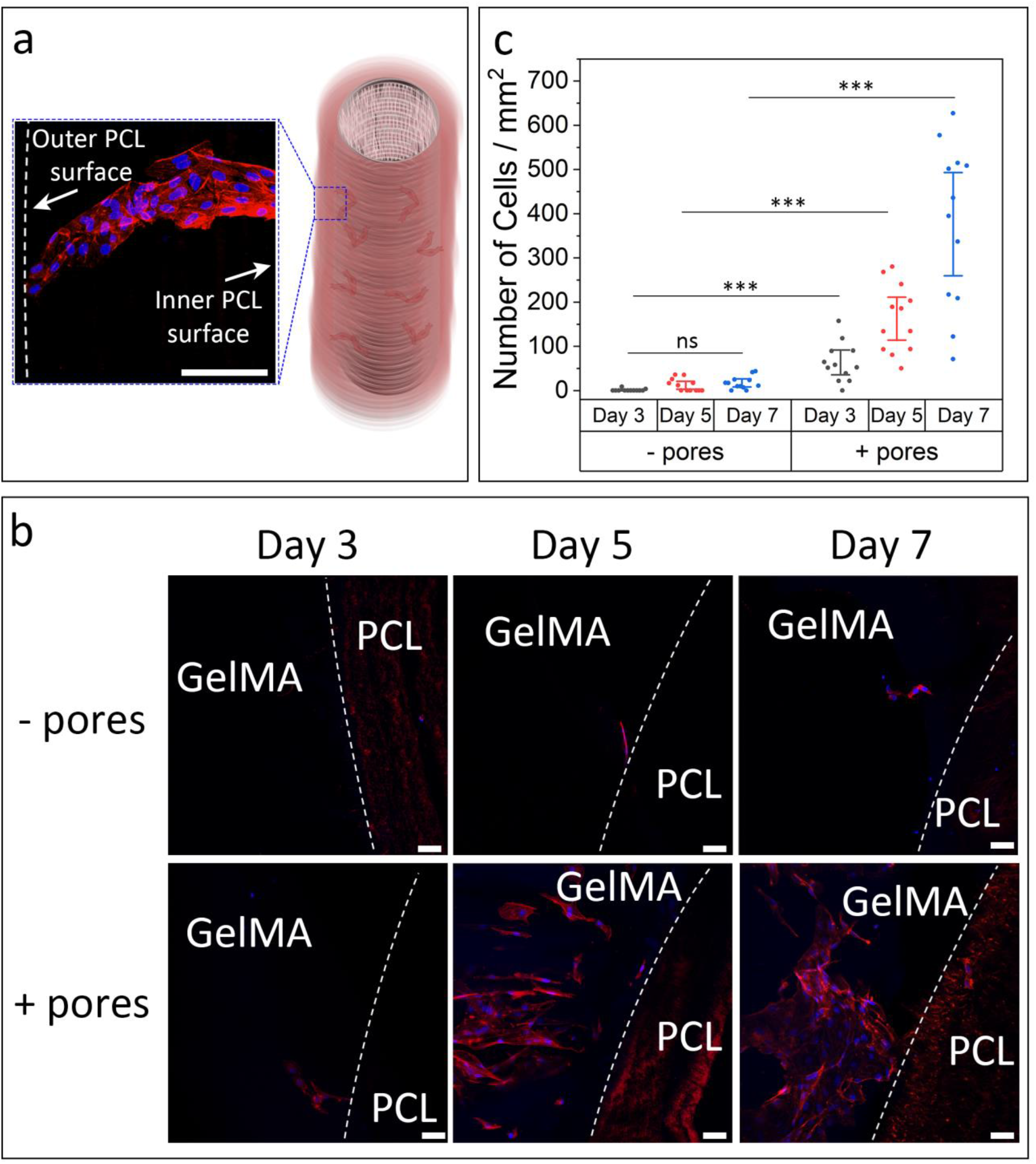
Cell migration from the PCL lumen toward the GelMA layer as a function of time after seeding the ECs. a) A sketch and a confocal image showing the migration of ECs from the lumen toward the surrounding GelMA layer through the PCL layer. b) Confocal images of ECs at the PCL/GelMA interface in samples with and without macropores; the dotted line indicates the edge of the PCL layer. c) Quantitative analysis of cell migration demonstrating the effect of macropores and culture time on the number of cells in the GelMA layer adjacent to the PCL layer. red: actin filaments stain (ActinRed™ 555), blue: nuclei stain (DAPI). Error bars represent the standard deviation (n = 12; ***: p < 0.001; ns: no significant difference, Student’s t-test). Scale bar = 50 μm.

### 2.4. Soft Hydrogels Promote Vascular Network Formation

**Figure 5a** illustrates the formation of a vascular network in the GelMA layer of the engineered tissue through EC migration from the lumen of the vascular graft. The density of a biodegradable hydrogel network regulates the ability of cells to migrate through the hydrogel, and the stiffness of the gel can impact the mechanobiology of embedded cells [36]. Thus, we assessed the ability of hydrogels with varying matrix densities to allow EC migration and vascular network formation. Specifically, we assessed these behaviors in 3, 7, and 10% w/v GelMA hydrogels. The Young’s modulus of the hydrogels increased with the concentration, as shown in our previous study: 4.3±0.9 kPa for 3% w/v, 43.7±5 kPa for 7% w/v, and 139±16 kPa for 10% w/v [19]. Interestingly, different migration and vascular network formation behavior was observed as a function of GelMA concentration, as seen in **Figure 5b**. Specifically, few ECs were found within the GelMA layers for the higher concentration 7 and 10% hydrogels at 7 days. Additionally, minimal to no vascular network formation was observed. However, a large number of ECs were found at the outer surface of the hydrogels. These results indicate that ECs migrated from the lumen to the edge of the GelMA and formed a monolayer on the external surface. Similar results were observed after 14 days, though there does seem to be some vascular network formation near the outer edge of the 7% hydrogel at the end point of the experiment. The difference in the vascularization behavior of ECs in the 7% and 10% w/v GelMA might be attributed to the difference in the polymer density. Several studies have shown that cells can degrade hydrogels with lower concentrations faster, which could explain the invasion of ECs after 14 days of culture in 7% w/v GelMA, but not in 10% w/v GelMA [37, 38]. In contrast, the ECs in 3% w/v GelMA formed a vascular network in the GelMA layer after 7 days of culture and the network density was increased at day 14. These results support our hypothesis that a hydrogel permeated with a vascular network can be created via migration of ECs through engineered macropores in a TEVG.

**Figure 5.**
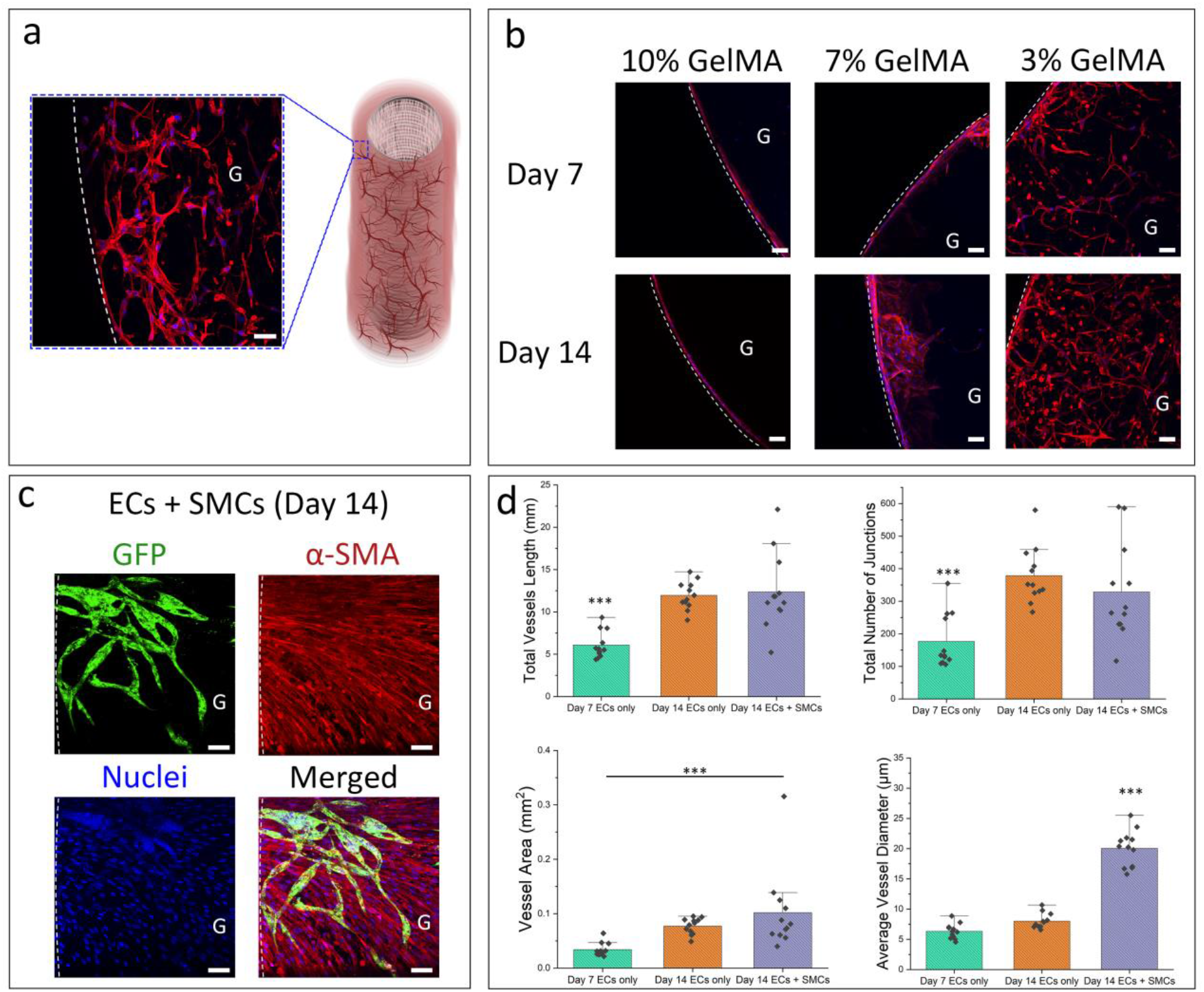
Vascular network formation in the GelMA layer. a) A schematic diagram and a confocal image showing a vascular network formed in the GelMA layer (3% w/v, G indicates the GelMA layer; dashed line indicates the outer edge of the sample, red: actin filaments stain (ActinRed™ 555), blue: nuclei stain (DAPI). b) Confocal images showing the vascular network formation in GelMA of different concentrations at different time points (red: actin filaments stain (ActinRed™ 555), blue: nuclei stain (DAPI). c) Confocal images showing ECs forming vascular networks in 3% w/v GelMA in the presence of SMCs (green: GFP-expressing ECs, red: α-SMA (Alexa Fluor 594), blue: Nuclei (DAPI)). d) Quantitative analysis of the vascular networks in 3% w/v GelMA. Total vessel length, number of junctions, and vessel area were determined using AngioTool, while ImageJ was used to measure average vessel diameter. Error bars represent the standard deviation (n = 12; ***: p < 0.001, one-way ANOVA). Scale bar = 50 μm.

To function as an engineered tissue, other cells beyond the ECs must be able to cohabitate in the hydrogel without impeding the formation of the vascular network. To assess the effect of co-culturing, EC network development was assessed in hydrogels that were pre-seeded with SMCs. **Figure 5c** illustrates that GFP-expressing ECs in 3% w/v GelMA formed a vascular network within the SMC-laden hydrogel after 14 days of culture. The confocal images were analyzed to quantify total vessel length, total number of junctions, vessel area, and average vessel diameter. Quantitative analysis of the vascular networks (no SMCs) showed that both the total vessel length and total number of junctions increased significantly from day 7 to 14 (p < 0.001), but no significant difference was observed between the samples with and without SMCs at day 14 (p = 0.325), as shown in **Figure 5d**. The analysis also showed that the vessel area was significantly larger between day 7 (0.03±0.01 mm^2^) and day 14 in the presence of the SMCs (0.07±0.01 mm^2^), but not without SMCs (0.1±0.07 mm^2^). The average vessel diameter was similar between day 7 (6.3±1.1 µm) and 14 without SMCs (7.9±1.2 µm) but increased significantly in the presence of SMCs (20±2.8 µm). This increase in vessel diameter likely reflects the maturing and stabilizing effect SMCs on the vascular network. This aligns with previous studies demonstrating that EC-SMC interactions promote vessel stability and size [39, 40]. These results illustrate that ECs can form vascular networks in the hydrogels in the presence of co-cultured cells and that the presence of SMCs may impact the vessel maturation. Furthermore, as the PCL degrades over time, the presence of circumferentially oriented SMCs would facilitate the formation of an organized medial layer, supporting the construct’s mechanical integrity.

### 2.5. Larger Engineered Tissues can be produced by incorporating multiple Tissue-Engineered Blood Vessels

We increased the size of the engineered tissue by fabricating a construct containing two TEVGs, seeded with ECs as described for constructs containing a single TEVG. The double-graft model was thicker, at 10 mm diameter, compared to the 4 mm diameter tissue described above. Based on the observation that the ECs migrated from the lumen of the PCL tube to the surrounding GelMA layer and formed a vascular network, we anticipated that the ECs would migrate from the lumens of both tubes and form a more extensive network in the GelMA layer, especially in the region between the tubes. **Figure 6a** shows confocal images of the engineered tissues after 14 days of culture. We observed endothelialized lumens, indicating that the endothelialization was not adversely affected by the increase in the tissue size. Additionally, the ECs formed a dense vascular network in the GelMA, with a higher density network observed in the regions between the tubes. Using light-sheet microscopy, we obtained images of the whole structure. **Figure 6b** shows various patterns of EC network formation, including migration from the vascular grafts, and network formation from the PCL tube side and from the GelMA side, as indicated by the arrows. These results suggest that the vascular network formation was not impaired by the increase in the tissue size. This engineered tissue is 1 cm in diameter, contains an embedded and three-dimensional vascular network, and is connected to endothelialized macrovessels that have the potential be sutured to NBVs, representing a significant advancement toward creating large and vascularized engineered tissues that can survive once transplanted *in vivo*. This vascularized tissue incorporating two central, medium sized vessels enhances the possibility that this tissue could be anastomosed to both arterial and venous circulations at an *in vivo* implant site. Future work will explore how effectively the vascular networks from each graft connect with each other, their survival and performance *in vivo*.

**Figure 6.**
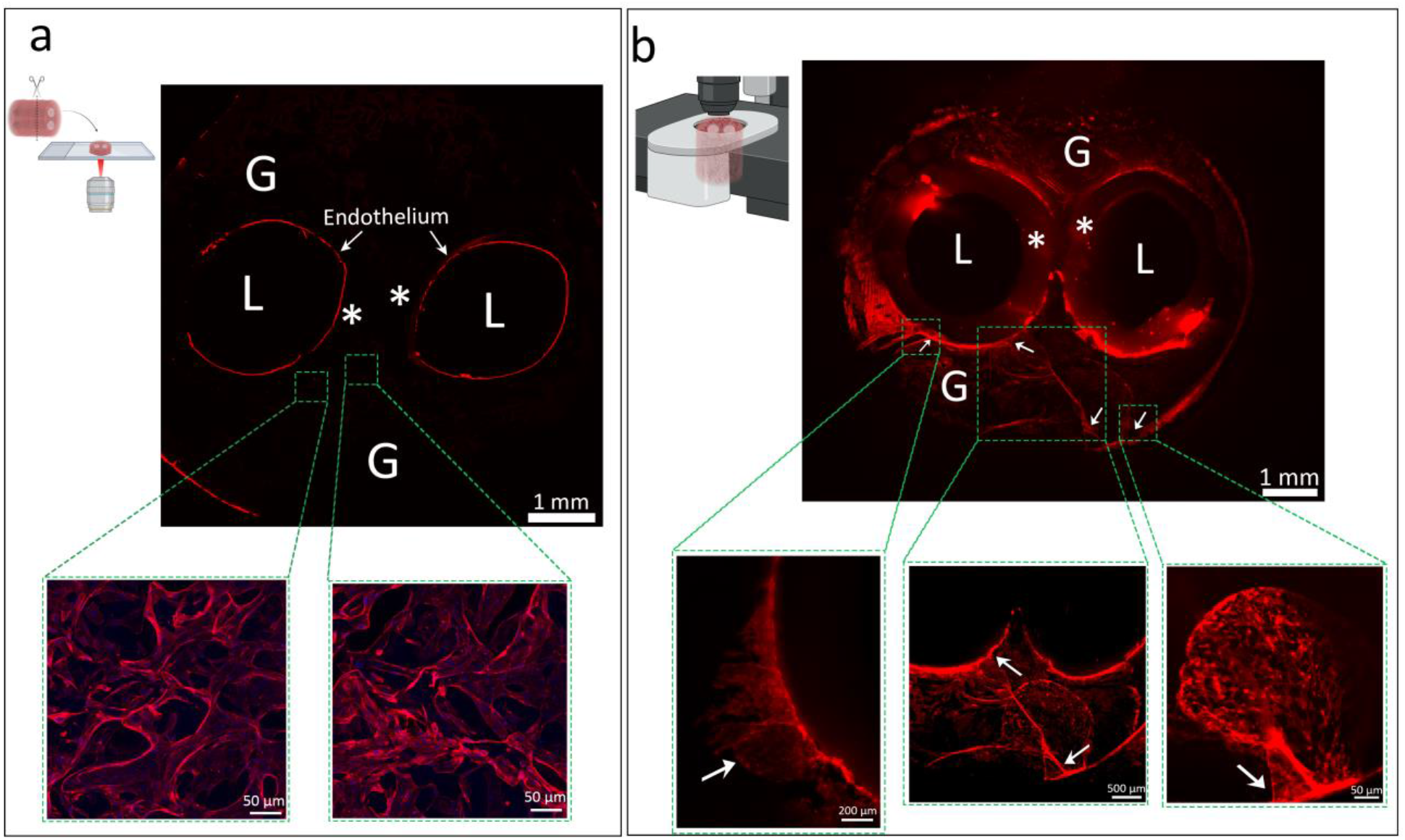
Vascularization of a large (1 cm dia.) tissue after 14 days of culture. a) Confocal images of the large tissue showing the ECs covering the lumen and forming an intensive network in the surrounding GelMA layer (red: actin filaments stain (ActinRed™ 555), blue: nuclei stain (DAPI). b) Lightsheet microscope images showing the overall vascular network in the GelMA layer with magnified images of selected regions showing the migration of cells and the vascular network formed; red: actin filaments stain (ActinRed™ 555). L indicates the lumen position, * indicates the PCL layer position and G indicates the GelMA layer position, the arrows indicate some vascular network formation instances.

## 3. Conclusion

The translation of vascularized tissues from the laboratory to the clinic is essential for the advancement of tissue engineering. However, this remains a challenge due to the weak mechanical properties of hydrogels and the impracticality of implanting conventional microfluidic systems. In this study, we developed a vascularized tissue surrounding a suturable TEVG to obtain a mechanically robust structure that has the potential for transplantation. The vascular graft was seeded with ECs that formed an endothelium after 5 days of culture. By introducing pores in the vascular graft, the ECs migrated from the lumen into the surrounding hydrogel and the migration rate increased after 7 days of culture as the endothelium reached confluence. Using low concentration GelMA (3%), the ECs formed a vascular network with and without the presence of SMCs. The tissue size can be increased by incorporating two vascular grafts, which resulted in a dense vascular network formation in a larger 1 cm diameter construct. This represents significant progress toward the translation of tissue-engineered constructs by unlocking the potential of solving the longstanding vascularization challenge that has limited the field for decades.

## 4. Materials and Methods

### 4.1. Electrospinning

The TEVGs were fabricated as described previously [19]. Briefly, a 12% w/v PCL (Mw 80 000 Da, Sigma-Aldrich, Germany) solution was electrospun onto a rotating collector rod (2.4 mm diameter, ER316L TIG filler wire, Hampdon, Australia) with a 12 kV voltage. Prior to cell seeding, the tubes were disinfected with ethanol and UV light, rinsed with phosphate-buffered saline (PBS, Gibco, USA), and incubated in complete EC growth medium (Lonza, USA).

### 4.2. Synthesis of Gelatin Methacryloyl (GelMA)

The synthesis of GelMA was reported previously [19]. In summary, a 10% gelatin type A solution in PBS was heated at 50°C for an hour, then methacrylic anhydride was slowly added (6% volume ratio) and reacted at 50°C for another hour. Quenching with PBS (2:1 volume ratio) and centrifugation separated the GelMA, which was then dialyzed in water for a week and lyophilized for storage. The degree of methacrylation was assessed by ^1^H NMR, comparing the spectra of GelMA and gelatin dissolved in D2O. Integration of specific proton signals revealed the extent of modification.

### 4.3. Micro-machining of the Electrospun Tubes

The PCL tube fabrication involved a micro-needle array stamping technique to create large pores through the tube walls. The array had 140 evenly spaced needles, each with a diameter of 300 µm, a length of 3 mm, and a spacing of 1.5 mm. The electrospun PCL tubes were mounted on a dental wax board (Proscitech, Australia) and pierced by pushing the micro-needle array through both walls of the tubes.

### 4.4. Increasing the Hydrophilicity of PCL

To enhance the hydrophilicity of PCL fibers and promote EC adhesion on the inner surface of electrospun PCL tubes, we followed a previously reported method [41]. We submerged PCL tubes in 1 M sodium hydroxide solution (Chem-Supply, Australia) for 4 h, rinsed them with water five times, and soaked them in water overnight.

### 4.5. Scanning Electron Microscopy

The pores in the electrospun PCL fibers were characterized by scanning electron microscopy (SEM) at an accelerating voltage of 10 kV (FlexSEM 1000, Hitachi, Japan). To prepare the PCL tubes, the samples were cut into 5 mm segments with a razor blade. The side view of the vascular grafts was examined by SEM. The samples were not sputter coated due to the low vacuum technology of the Hitachi SEM microscope that eliminates the need for sputter coating, enabling direct observation of non-conductive specimen surfaces [42].

### 4.6. Preparation of GelMA Mixtures

The GelMA solution was prepared by dissolving GelMA (3, 7, or 10% w/v) in EC media at 37 °C for 60 min in an incubator. Then, the photoinitiator lithium phenyl-2,4,6-trimethylbenzoylphosphinate (LAP, 0.06% w/v, Sigma-Aldrich, Germany) and gelatin type A (1.6% w/v) were added to the solution. Gelatin was added to the mixture to temporarily enhance the cast layer’s structural integrity after photocrosslinking, facilitating smoother removal from the mold. The solution was then sterilized by passing through a 0.22 μm PES syringe filter (Merck, Germany).

### 4.7. Fabrication of Suturable Vascularized Constructs

The fabrication process for the suturable vascularization models is illustrated in Figure 1b. A sterile polytetrafluoroethylene-coated rod (diameter: 2.5 mm) was inserted into the lumen of the PCL tube and placed inside a transparent polypropylene mold (4.4 mm inner diameter for the single vessel tissue, or 10.5 mm inner diameter for the double-vessel tissue). The mold was then filled with the preheated GelMA mixture (with or without SMCs at a density of 500 000 cells/mL) until it was fully submerged. The rod prevented the GelMA mixture entering the lumen. The mold was cooled in an ice-bath for 1 min to induce physical gelation of the GelMA mixture, followed by UV irradiation for 60 s (OmniCure S1500, 320−500 nm filter, 30 mW/cm^2^) to achieve photo-crosslinking. The mold was then immersed in the ice-bath for another 3 min before the bilayered GelMA/PCL construct was released from the mold and the rod.

### 4.8. Mechanical Testing of the Vascular Grafts

To measure the circumferential ultimate tensile strength (UTS) and Young’s modulus of the vascular grafts, 5 mm segments were cut and secured to stainless-steel holders on an Instron 5944 Microtester with a 50 N load cell. The segments were stretched at a rate of 50 mm/min according to ISO 7198:2017. A burst pressure test was performed by connecting a 2.5 cm segment of vascular graft to a pressure transducer (Lutron, USA) and a pressure meter (Sper scientific, USA). The graft was filled with Vaseline (Unilever, USA) using a syringe pump (Harvard Apparatus, USA) at a flow rate of 1 mL/min until rupture. The maximum pressure exerted during this process was then recorded. Suture retention testing was conducted following the ISO 7198:2017 protocol. Briefly, 1 cm segments of vascular graft were sutured (Ethicon 5−0, Scotland) 2 mm from the edge, creating a half loop on one side. The base of the graft was secured in mechanical grips, and the suture was pulled at a speed of 50 mm/min using an Instron 5944 Microtester until the wall failed. Young’s modulus was obtained from the linear region of the stress/strain curve.

### 4.9. Cell Culture

Human umbilical vein ECs (Lonza, USA) and GFP expressing human umbilical vein ECs (Angio-Proteomie, USA) were cultured in EGM-2 medium (Lonza, USA) and endothelial growth medium ECs (Angio-Proteomie, USA), respectively, with 1% v/v antibiotic-antimycotic solution (Gibco, USA). They were seeded into the vascular graft at passage 5 using a micropipette. Human coronary artery SMCs (Lifeline, USA) were cultured in SMC Medium (Lifeline, USA) with 1% v/v antibiotic-antimycotic solution and encapsulated in GelMA during passage 5. Cells were grown in 75 cm^2^ flasks (Corning, USA) and the medium was changed every two days until 70% confluence. ECs were seeded (100 µL, 3 × 10^6^ cells/mL) onto the lumen of the TEVG scaffolds. Both ends of the scaffolds were subsequently sealed, and the constructs were subjected to gentle rotational (2 rpm) within a custom-designed bioreactor for 4 h in an incubator. This rotational regime promoted uniform cell attachment throughout the inner luminal area. Subsequently, the EC-seeded TEVGs were transferred to static culture flasks for further analyses.

### 4.10. Immunofluorescence Analysis

The vascular grafts were transferred to a 15 mL tube after the cultivation period and washed with PBS three times. The grafts were fixed in 4% paraformaldehyde (Scharlau, Spain) at 4 °C for 4 h and then sliced into 5 mm segments using a razor blade. The segments of single and double vessel tissue were placed in 96-well and 48-well plates, respectively, for staining. The segments were washed with PBS three times and permeabilized with 0.1% Triton X-100 (Labchem, USA). ActinRed 555 ReadyProbes Reagent (Thermo Fisher, USA) was used to stain the actin filaments for 90 min. The nuclei were stained with a 1:10,000 dilution of DAPI (Sigma-Aldrich, Germany) after washing the segments with PBS three times. For CD31 and αSMA staining, the segments were blocked with 1% albumin bovine serum (Sigma-Aldrich, Germany) for one hour and then incubated with Alexa Fluor 488 anti-CD31 antibody and anti-alpha smooth muscle actin antibody (abcam, United Kingdom) overnight. The next day, donkey antirabbit IgG H&L (Alexa Fluor 594) was added for 4 h. The segments were stored in PBS at 4 °C until imaging. Confocal images were acquired using a Nikon A1R+ Confocal Microscope (Nikon, Japan). To minimize light scattering and enhance depth penetration during lightsheet imaging, the samples were immersed in a 30% v/v glycerol (Thermo Fisher, USA) solution in distilled water, eliminating the need for separate clearing steps due to the inherent transparency of GelMA. Lightsheet images were obtained with an UltraMicroscope II (Miltenyi Biotec, Germany).

### 4.11. Evaluation of Endothelial Coverage

EC coverage was quantified by confocal microscopy images of randomly selected regions of the inner lumen surface. The red channel containing the actin filaments staining (ActinRed™ 555) of ECs was extracted from the images and binarized using the threshold function in ImageJ. The black pixels, corresponding to the cells, were measured as a percentage of the total area using the measurement function in ImageJ.

### 4.12. Evaluation of Cell Migration

The migration of ECs was assessed by confocal microscopy. Samples were randomly selected and imaged at the PCL/GelMA interface of samples with and without macropores. The cell nuclei were stained and the number of cells in the GelMA region was determined using ImageJ software with the cell counter plugin. The cell density was calculated by dividing the cell number by the area of GelMA in each image. For migrating cell density, only cells within 200 µm of the PCL layer were counted.

### 4.13. Evaluation of Vascular Network

The vascular network was quantified by confocal microscopy at day 7 and at day 14 with and without SMCs. Samples were randomly selected, and images taken of the GelMA layer. The actin filaments were stained, and the resulting images were analyzed using AngioTool software [43], an open-source image analysis platform specifically designed for quantifying of angiogenesis (ImageJ plugin version 0.6a, National Institutes of Health, USA). AngioTool was used to measure total vessel length, total number of junctions, and vessel area.Average vessel diameter was determined using ImageJ software. Evenly spaced horizontal grid lines (approximately 9 per image) were randomly drawn by ImageJ and the diameter of 20 vessels intersecting with the lines was calculated per image.

### 4.14. Statistical Analysis

Statistical analysis was performed using Minitab 21.4.0 software. To compare the means of two data sets, Student t tests were applied. To compare the means of more than two data sets, one-way ANOVA with a Tukey’s post hoc test was used. The level of significance was set at p < 0.05.

## 5. Acknowledgements

This work was partially funded through D.H.’s Australian Research Council Future Fellowship (FT190100280). This work was performed in part at the Materials Characterisation and Fabrication Platform (MCFP) at the University of Melbourne and the Victorian Node of the Australian National Fabrication Facility (ANFF). Furthermore, we would like to gratefully acknowledge the Biological Optical Microscopy Platform for their support and assistance with confocal microscopy imaging in this work. H.A. gratefully acknowledge the support of the University of Melbourne and an Australian Government Research Training Program Scholarship (Melbourne International Research Scholarship). A.O. gratefully acknowledges support of the Australian Research Council, The Brenda Shanahan Charitable Foundation, and St Vincent’s Hospital Melbourne.

## 7. Data Availability

Data will be made available on request.

