## Supplemental information for "Hierarchically Vascularized and Implantable Tissue Constructs created through Angiogenesis from Tissue-Engineered Vascular Grafts"

### Supporting Information

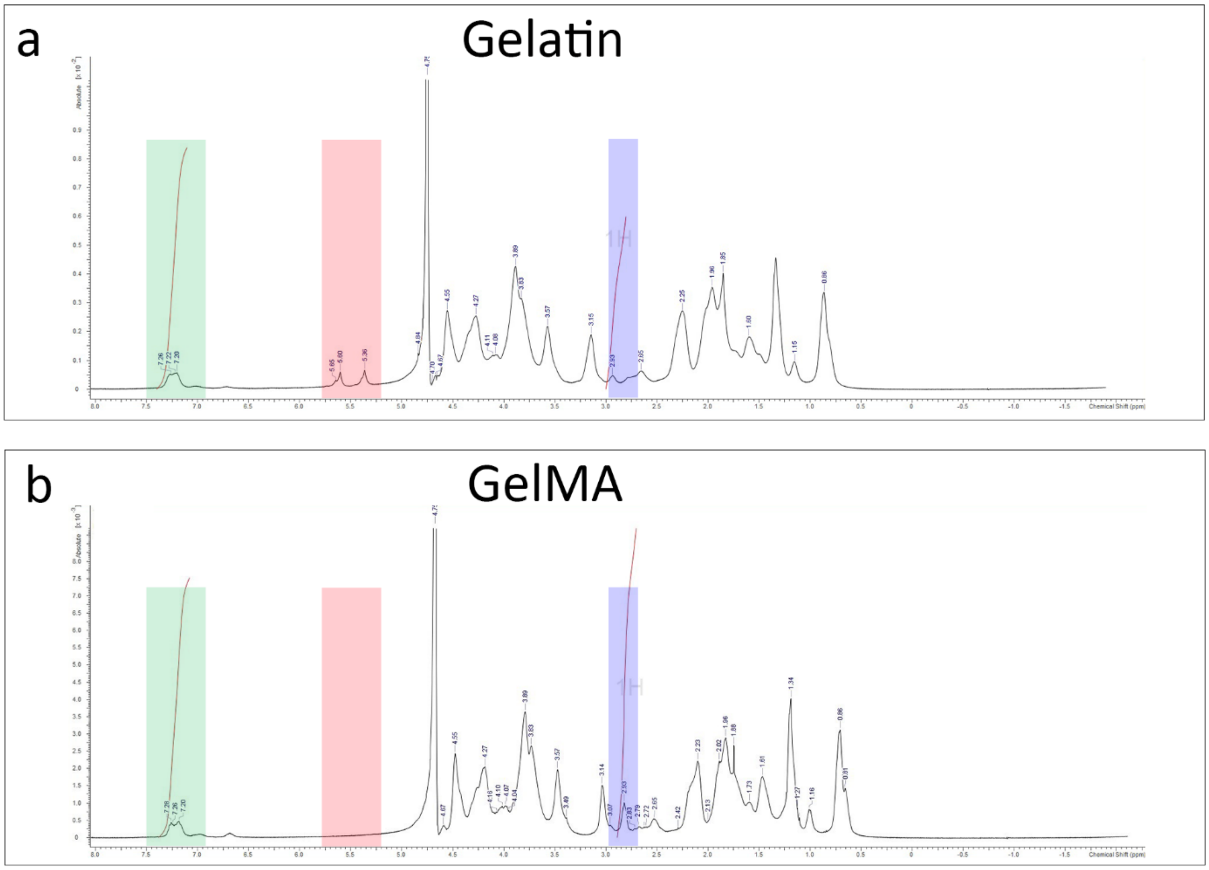


Figure S1. $^{1}$H NMR spectra of gelatin and GelMA that were used to quantify the degree of methacrylation. a) $^{1}$H NMR spectrum of unmodified 5% w/v gelatin. b) NMR spectrum of 5% w/v gelatin-methacryloyl (GelMA). Green highlight: Aromatic amino acid. Red highlight: Methacrylate vinyl group of the MA. Blue highlight: Lysine methylene.
